# Sex and interferon gamma signaling regulate microglia migration in adult mouse cortex *in vivo*

**DOI:** 10.1101/2023.02.02.526857

**Authors:** Roobina Boghozian, Sorabh Sharma, Manjinder Cheema, Craig E. Brown

## Abstract

Although microglia possess the unique ability to migrate, whether mobility is evident in all microglia, is sex dependent, and what molecular mechanisms drive this, is poorly understood in the adult brain. Using longitudinal *in vivo* imaging of sparsely labelled microglia, we find a relatively small population of microglia are mobile under normal conditions. Following injury (microbleed), the population of mobile microglia increased in a sex-dependent manner, with male microglia migrating significantly greater distances towards the microbleed relative to their female counterparts. To understand the signaling pathways involved, we interrogated the role of interferon gamma (IFNγ). Our data show that in male mice, stimulating microglia with IFNγ or inducible microglial specific knockdown of IFNγ receptor 1 stimulates or inhibits migration, respectively, whereas female microglia were generally unaffected. These findings highlight the diversity of microglia migratory responses to injury, its dependence on sex and the signaling mechanisms that modulate this behavior.

## Introduction

Microglia are innate immune cells that reside within the brain^1^. Seminal imaging studies revealed the highly dynamic behavior of these cells^2,3^. Under normal conditions, microglia continually re-arrange their processes to scan the local microenvironment, a behavior known as surveillance^4,5^. Following injury, microglial processes grow towards and engulf cellular debris, referred to as process chemotaxis^6,7^. However, microglia also possess the capacity for movements of the soma or migration^8^. In early development, microglia derived from the yolk sack migrate into the brain^9^, usually along nascent brain vasculature to populate their respective territories^10,11^. As the brain matures, there is some ambiguity regarding the extent to which microglia mobilize under normal/homeostatic conditions. For example, some longitudinal imaging studies report that microglia are very long-lived (median life-time over 15 months) and stable in their somatic position during adulthood^10,12–14^. Other studies using fluorescent fate mapping or imaging have shown that a fraction of cells appeared in new positons. These changes in position were largely attributed to a balanced combination of cell proliferation and death^12,15–17^. Some instances of cell body movements were noted although on a very limited scale (≤15μm displacement)^18,19^.

The mobility of microglia towards sites of injury or toxic proteins in neurodegenerative disease, could be important for mollifying toxicity and resolving injury ^20–23^ but could also contribute to the spread of toxic proteins (eg. Amyloid beta)^24^ and pruning of synapses^25–27^. Indeed, microglia migration has been noted by their clustered appearance around sites of injury^15,28,29^. For example, Cx3Cr1-eGFP labelled microglia/macrophages show directed movements towards a hemorrhage in the first 48h^14^ or millimeters away from an ischemic stroke over weeks^30^. Although these studies have been very informative, they have relied on transgenic mice that express a fluorescent reporter in most, if not all microglia. Thus, unambiguously tracking individual cell movements was nearly impossible, especially if a cell were to traverse a long distance in a short period of time, or if cells were to cluster in one spot, which invariably occurs after focal injury. Furthermore, since the transgenic mice used in these studies also label brain resident macrophages or infiltrative leukocytes^44^, attributing movements specifically to microglia was not conclusive. Thus, there has been little direct, quantitative tracking of migratory behaviours in individual microglia after injury.

Although we have a better understanding of the molecular mechanisms that regulate microglial surveillance and process chemotaxis^31,32^, the pathways that govern migration, beyond the role of P2RY12 signaling^16^, has been mostly limited to *in vitro* studies or retina^33^. Pro-inflammatory cytokines such as IFNγ, are a likely candidate given that IFNγ receptors are expressed on microglia and its signaling is induced by brain injury such as stroke or hemorrhage, where it regulates many facets of macrophage behavior^34,35^. In cell or slice cultures, IFNγ exposure triggers inflammatory cytokine expression, phagocytic activity and increases mobility^36,37^. Interestingly, cultured microglial responses to IFNγ can differ by sex^38^, where male cells exhibit greater motility in the trans-well assay than their female counterparts. Sex differences also exist in microglia responses to ischemic stroke and/or pro-inflammatory stimuli^39–43^. Based on these studies, it stands to reason that IFNγ signaling and sex could regulate migratory behaviors of microglia in the mature brain.

Despite our increased understanding of microglial biology, important questions remain concerning microglia mobility in the mature mammalian brain. For example, what fraction of microglia are mobile after vascular injury and are all cells equally adept at moving? How far can these cells travel per day? Is mobility affected by sex? What molecular mechanisms regulate mobility in the mature brain? In the present study we tracked the mobility of individual cells using an inducible, microglia specific cre-driver mouse. Our experiments show that while the population of mobile microglia is quite small under normal conditions, it can be rapidly expanded following injury. Further we reveal that that mobility was strongly influenced by biological sex and IFNγ signaling. Collectively, these results provide new information about the factors (injury, sex, cytokine signaling) that regulate microglia mobilization in the mature mammalian brain.

## Results

### Methodological considerations for tracking mobile microglia *in vivo*

For the sake of clarity, the terms microglial “mobility or movements” herein refers to the movements of microglia cell bodies to new positions, not movement of processes. To quantify microglial mobility *in vivo*, we had to first define what distance would constitute a true movement. Cells within the living brain can be pushed or displaced small distances for a number of reasons (changes in large vessel diameter, natural contortion of the brain) and repeated 3-dimensional measurements possess an intrinsic degree of error. Therefore we measured the distance between GFP labelled VIP interneurons and a fiducial landmark at 12 hour (h) intervals in mice lightly anesthetized with isoflurane (**Supp. Fig. 1**). Since fully differentiated neurons in the adult mouse cortex do not migrate, this allowed us to estimate measurement error *in vivo* and thus identify true migration. By tracking 206 neurons in 3 mice, we found the average displacement of cells per 12h period was 1.84±1.45μm and 1.61±1.53μm (**Supp. Fig. 1**). To minimize the possibility of false positive detection of cell soma “movement”, we determined that any distance ≥7.46μm per 12h period (4 SD above mean) represented true cell movement. This threshold formed the basis for what we defined as a “mobile” cell.

To track the migration of microglia in adult mice, we first utilized heterozygous Cx3cr1-eGFP reporter mice^44^, as described in previous studies that reported somatic movements^16,18^. These mice were with implanted with a craniectomy based cranial window and were allowed to recover at least 5 weeks before imaging. We should note that thin skull imaging windows were also prepared although in our hands, this procedure often acutely disrupted superficial blood vessels and was associated with sub-optimal signal to noise for deeper imaging. Consistent with previous imaging studies^16,18^, we found that the majority of putative microglia remained in the same position over a 24h period (96.2% were “stable”, **Fig. 1A**). However, a small fraction of cells appeared to either move a short distance (see yellow arrow in **Fig. 1A**), appeared or disappeared (termed “unstable” microglia; move: 3.1% vs new: 0.59% vs lost: 0.24%; see yellow arrows in **Supp. Fig. 2A**). Unfortunately, because the density of cell labelling was high, the shape of cells changed constantly and both microglia and macrophages express eGFP; we could not be sure if cells that changed (say appeared or disappeared) were microglia that moved a long distance or perhaps were infiltrative macrophages. Furthermore, we could not track single cell movements after injury when cells rapidly cluster around a focal injury site (**Supp. Fig. 2B**).

**Figure 1.**
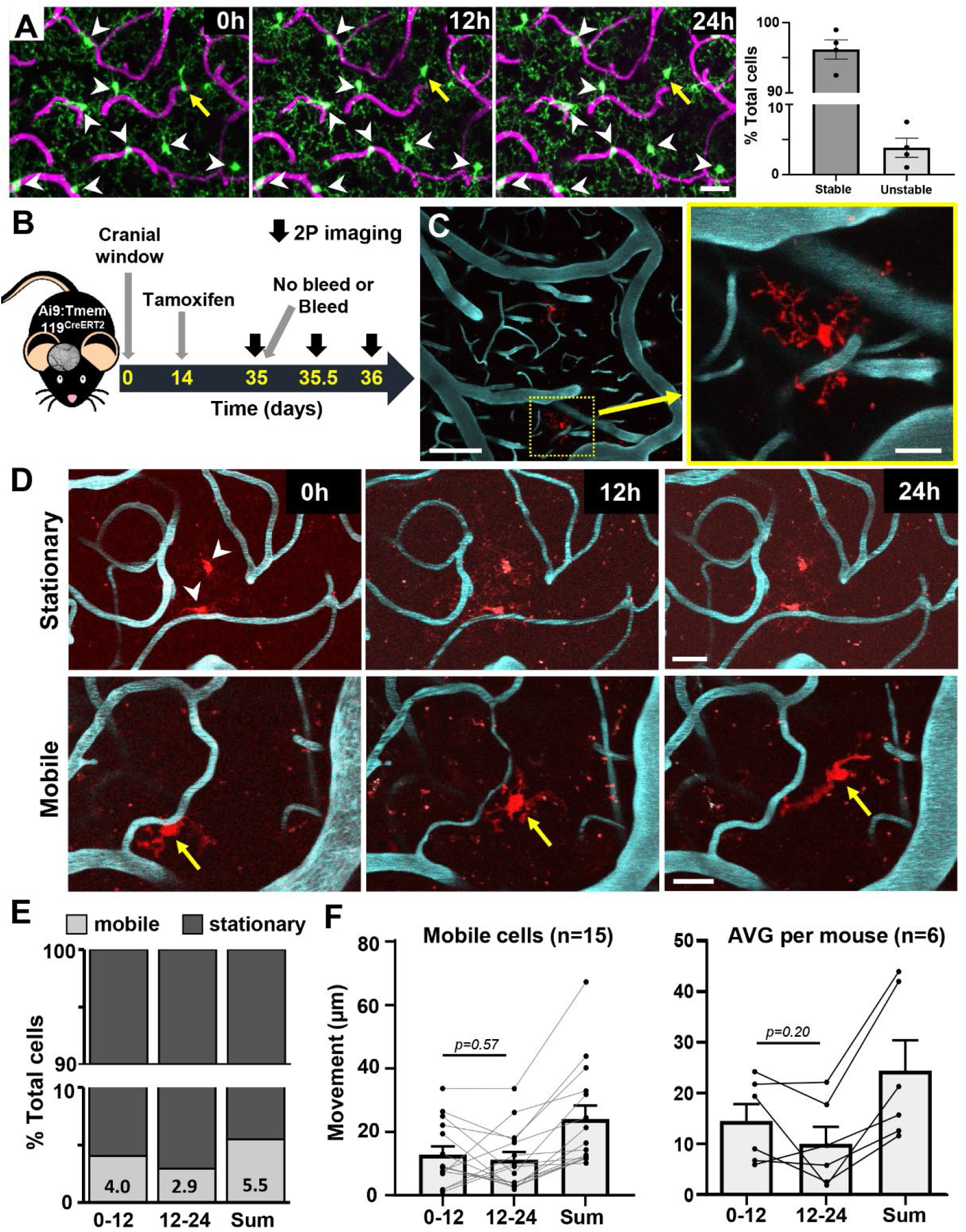
A small population of microglia are mobile in the normal adult brain. (A) Time lapse 2-photon maximum intensity projection images of Cx3cr1-eGFP labelled microglia/macrophages and vasculature over 12h intervals (left). The % cells that were “stable” or “unstable” (ie. those that moved, appeared or disappeared) is shown on the right. Scale bars, 20μm. (B) Schematic showing timeline of experimental procedures and imaging. (C) *In vivo* maximum intensity projection images showing sparse tdTomato labelled microglia and FITC labeled blood vessels in the somatosensory cortex. Scale bars, 100 and 20μm. (D) Time-lapse *in vivo* images of a stationary (top row) or mobile (bottom row) microglia over a 24h period. Scale bars, 20μm. (E) Fraction of mobile or stationary microglia (out of 272 cells, n=9 mice, 5 male and 4 female) imaged over 0-12h, 12-24h or the Sum of each window. Any microglia that moved >7.46μm in a 12h period was considered “mobile”. (F) Distance moved for each mobile microglia (left) or the average distance per mouse (right). Two-tailed paired t-test indicated no difference between 0-12h and 12-24h periods in distance travelled per cell (t_(14)_=0.59, p=0.57), or per mouse (t_(5)_=1.48, p=0.20). Data are mean ± SEM.

To track the movements of individual cells, we crossed the tamoxifen inducible and microglia specific Tmem119-CreERT2 mouse line with cre-dependent tdTomato reporter mice (Ai9). Consistent with its original description^45^, low dose injection of tamoxifen 3 weeks before the start of imaging yielded very sparse fluorescent labelling of microglia (2.84±2.9% of total microglia population, based on 52 areas in 16 mice) in the adult mouse brain (**Fig. 1B,C**). Furthermore, in the absence of tamoxifen, we did not find evidence of recombination (tdTomato labelled microglia), consistent with a recent study^46^.

### A small population of microglia are mobile under normal conditions

Having optimized methodologies to track microglia *in vivo*, we then imaged their mobility in 12h intervals over a 24h period (**Fig. 1B**). This interval was based on pilot work indicating that considerable movement was possible in a 12h period (especially after injury), thereby providing an interval of sufficient length to allow movement to occur, but not too long to cause us to lose track of individual cells. Under normal conditions, the majority of microglia were stationary (**Fig. 1D,E**; 94.5% of 272 cells over 24h), mirroring our estimate in Cx3cr1-eGFP mice. A small fraction of microglia clearly moved over this period, some upwards of 40-60μm (**Fig. 1D,E**; 5.5%, 15/272 cells). On average, microglia classified as “mobile” moved equivalent distances for each 12h period (~12.8 vs 11.2μm) for a mean of 24μm over 24h (**Fig. 1F**). We should note that since mice were imaged at the start of their light (8am; 0-12h) or dark cycle (8pm, 12-24h), we did not detect a circadian difference in movement per cell (**Fig. 1F** left panel, 2 tailed t-test, t_(14)_=0.59, p=0.57) or per mouse (**Fig. 1F** right panel, 2 tailed t-test, t_(5)_=1.48, p=0.20). Stratifying the movement of these mobile cells based on sex, did not reveal a significant effect of time or sex (2-way ANOVA, Time: F(1,26)=0.39, p=0.54, Sex: F(1,26)=4.09, p=0.053), although there was a trend towards reduced mobility in female cells (sum of movement over 24h for males 28.94±18.1μm vs 14.3±4.03μm in females). These results indicate that a small population of microglia are actively moving about in the uninjured brain.

### Sex dictates microglial mobility in response to vascular injury

In order to stimulate microglia mobility, we assessed responses to a laser induced cerebral microbleed (“CMB”; vessels 3.5-6μm in diameter, see **Fig 1B** for experimental outline). *In vivo* imaging showed that some microglia migrated long distances towards the bleed over a 24h period (see example in **Fig. 2A**). As we did not find a significant effect of sex on mobility under normal conditions, we first analyzed mobility with data from both sexes pooled (see **Fig. 2B-D**). When compared to uninjured controls, the induction of a CMB significantly increased the fraction of mobile microglia over 24h (**Fig. 2B**, χ2=94.5, p<0.01). However to our surprise, the absolute distance of migration (**Fig. 2C**) and the net movement of mobile microglia towards the CMB (**Fig. 2D**), did not change significantly when compared to un-injured controls.

**Figure 2.**
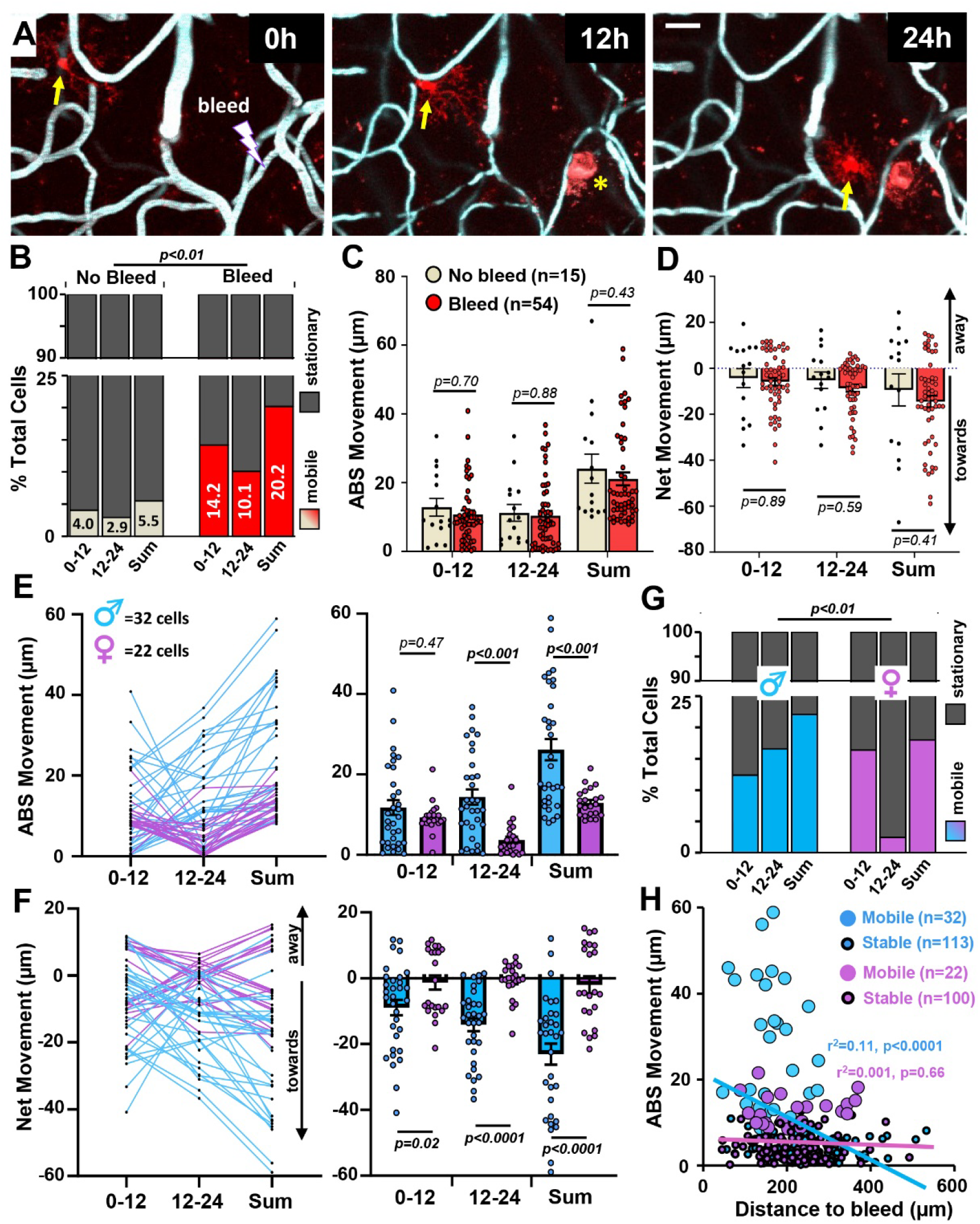
Sex specific differences in microglial mobility after focal vascular injury. (A) *In vivo* two photon images showing a microglia (yellow arrow) in a male mouse migrate towards a micro-bleed over 24h period. Note that vessel ablation leads to red emitting auto-fluoresce at the injury site (yellow asterisks). Scale bar, 20μm. (B) Fraction of mobile and stationary microglia imaged over a 24h period under normal (no bleed; 272 cells from 9 mice) or Bleed (267 cells from 13 mice) conditions, with data from both sexes pooled together. Chi-square analysis indicates the fraction of mobile microglia increased significantly after injury (χ2 =109.9, p<0.01). (C) Bar graphs show the absolute distance travelled by mobile microglia in No Bleed (15 cells) and Bleed (54 cells) conditions for each 12h interval or the sum over 24h, with data from both sexes pooled together. Two-tailed unpaired t-tests were used to calculate p values. Data are mean ± SEM. (D) The net distance travelled by mobile microglia towards or away from the injury site (or an equivalent landmark in controls), with data from both sexes pooled together. Two-tailed unpaired t-tests were used to calculate p values. Data are mean ± SEM. (E) Graphs show the absolute movement of each mobile microglia plotted over time (left) or the group average (right) after injury, stratified by sex (32 cells from 7 males and 22 cells from 6 female mice). Two-tailed unpaired t-tests were used to calculate p values. Data are mean ± SEM. (F) The net movement of each mobile microglia plotted over time (left) or the group average (right) after injury, stratified by sex. Two-tailed unpaired t-tests to generate p values for comparisons between sexes. Data are mean ± SEM. (G) Fraction of mobile or stationary microglia for male and female mice after injury. Chi-square analysis indicates males had more mobile microglia after injury than female mice (χ2 =9.33, p<0.01). (H) Linear regression analysis in both male and female mice shows the relationship between how far a microglia moves over a 24h period as a function of each cell’s initial distance from the bleed. Note the moderate and significant relationship in male but not female mice. Large open circles represent mobile microglia while smaller dots with dark outline represent stationary microglia.

Since previous reports have shown that male microglia have a higher capacity to respond to ATP signals, display a more pro-inflammatory pattern of gene expression and can appear morphologically different than females^41–43^, we then stratified our data set by sex. This revealed that microglia from male mice migrated significantly greater absolute distances than female cells over the 24h period, primarily in the later 12-24h post-bleed time window (**Fig. 2E;** 2-way ANOVA, Sex: F_(1, 104)_=15.6, p=0.0001; Time: F_(1, 104)_=0.81, p=0.37; Sex x Time: F_(1, 104)_=6.02, p=0.02). Furthermore, microglia in male mice showed significantly greater net movement towards the CMB than female microglia, who displayed little directionality on average (**Fig. 2F;** 2-way ANOVA, Sex: F_(1,104)_=25.3, p<0.0001; Time: F_(1,104)_=1.21, p=0.27; Sex x Time: F_(1,104)_=1.99, p=0.16). The fraction of mobile microglia in male mice was significantly greater than in female mice (**Fig. 2G**; χ2=9.33, p<0.01). And finally, as it was plausible that a cells’ relative distance from the CMB could influence mobility, we found in male mice that microglia closer to the bleed were more likely to migrate (**Fig. 2H,** r^2^=0.11, p<0.0001), whereas in female mice, this relationship disappeared (**Fig. 2H,** r^2^= 0.01, p=0.37). We should note that the size of the bleed, based on the area of dye extravasation, was not significantly different between sexes (**Supp. Fig. 3;** t_(23.9)_=0.93, p=0.36). In addition to migratory behavior after injury (see examples **Supp. Fig. 4A,B),** we also noted possible examples of microglia proliferation (**Supp. Fig. 5A,B)**, although these were very infrequent events. Collectively, these results indicate that sex has a strong influence on mobility after vascular injury.

Given that sex and distance from injury were not the only variables at play in these experiments, we examined other factors that could influence microglia mobility. First, we considered a cell’s depth from the cortical surface. Parsing microglia based on whether they were stationary or mobile did not reveal any systematic difference in cortical depth (**Supp. Fig. 6A**), nor was there any significant relationship between movement distance and depth, even when stratifying cells by sex (**Supp. Fig. 6B**). Given there is growing appreciation for the diversity of microglia, especially those that reside along capillaries (referred to as capillary associated microglia or “CAM”) that modulate blood flow^47,48^, we assessed mobility patterns in these cells. Our analysis of CAM vs Non-CAM from both sexes did not show any significant differences in either the absolute or net movement of these cells after bleed (**Supp. Fig. 6C,D**). Separating our analysis based on sex did not reveal any significant differences in movement between CAM vs Non-CAM (**Supp. Fig. 6C,D**). In summary, our findings indicate that a cell’s depth below the cortical surface or whether it was associated with a capillary, does not significantly influence mobility.

### IFNγ signaling regulates mobility in a sex-specific manner

Our next goal was to understand the molecular mechanisms that regulate microglial mobility. Previous studies have shown that IFNγ signaling is associated with a pro-inflammatory/disease related gene signature^36,49^, modulates microglial process chemotaxis *in vivo*^50^; and preferentially stimulates migration in male microglia *in vitro*^38^. To this end, we employed different approaches for enhancing or blunting IFNγ signaling *in vivo*. To stimulate IFNγ signaling, we intravenously injected IFNγ or vehicle/control protein solution into Ai9:Tmem119cre (referred to as wild-type or “WT” controls) minutes before the induction of injury (**Fig. 3A**). We reasoned that IFNγ protein in the blood would easily leach into the brain parenchyma after vessel rupture and stimulate microglia. Furthermore, we could visually confirm the extravasation of fluorescent blood plasma after vessel injury (see **Supp. Fig. 3**). To knock down IFNγ signaling specifically within microglia, we crossed the Ai9:Tmem119cre line with IFNγ receptor 1 floxed mice (*Ifngr1*^fl/fl^), which previous studies have used to induce cell specific knockdown of IFNγ signaling^51–53^. This approach allowed us to simultaneously turn on a cre-dependent fluorescent reporter in a sparse population of microglia with cre-recombinase dependent knockdown of *ifngr1* (“*Ifngr1* KD”, see **Fig. 3B**).

**Figure 3.**
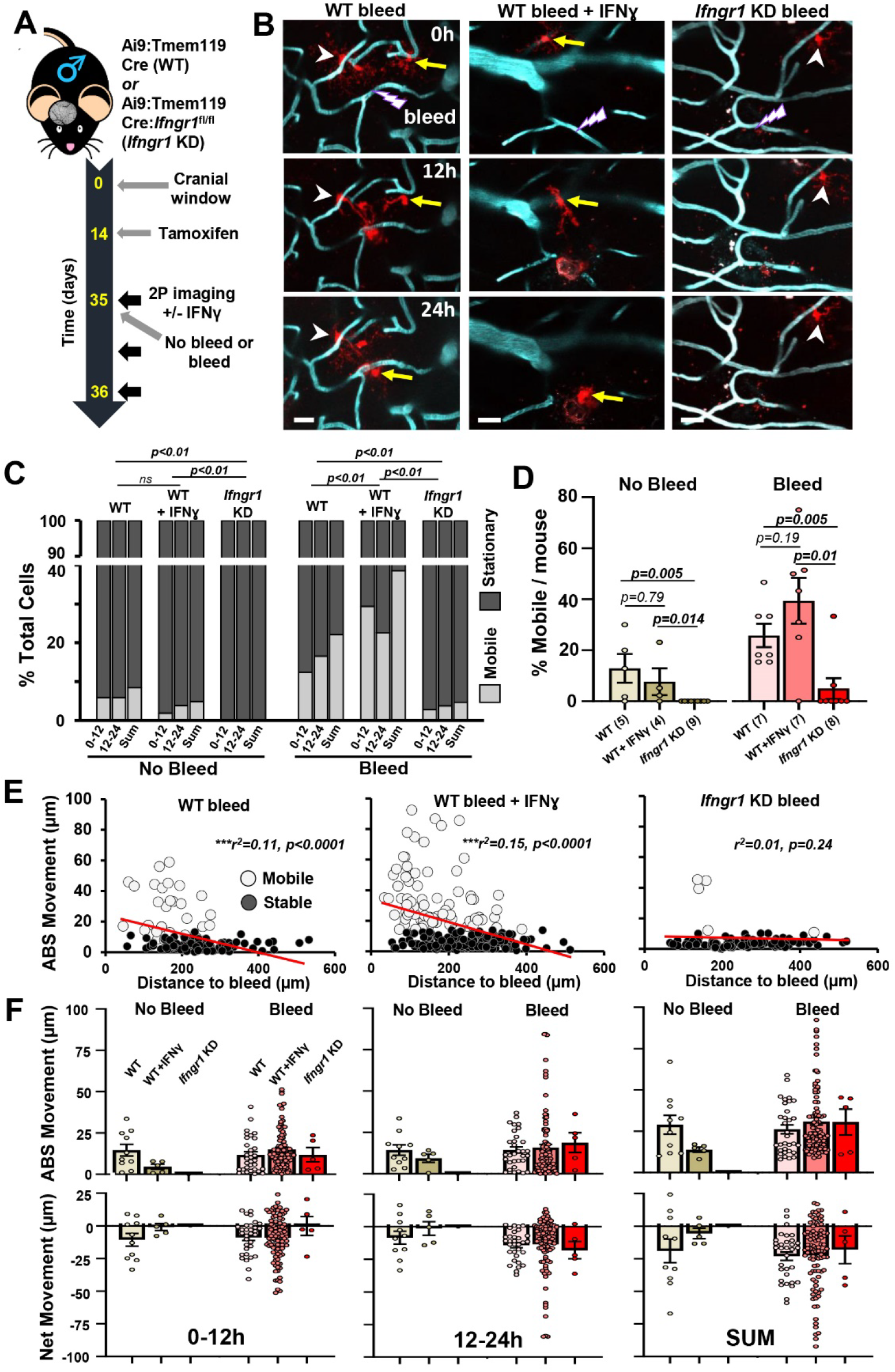
IFNγ signaling regulates microglial mobility after microbleed in male mice. (A) Schematic showing timeline of experimental procedures and imaging. All experiments and data shown in this figure originate from male mice. (B) *In vivo* two-photon images showing the movement of microglia after micro-bleed in wild-type (WT) controls, WT injected with IFNγ or microglia specific knockdown of *Ifngr1* (*Ifngr1* KD). Mobile and stationary microglia denoted by yellow arrow or white arrowhead, respectively. Scale bar, 20μm. (C) Fraction of mobile or stationary microglia under normal no bleed and bleed conditions for each of the three experimental groups (WT no bleed and bleed: 119 and 145 total cells; WT+IFNγ no bleed and bleed: 105 and 252 total cells; *Ifngr1* KD no bleed and bleed: 122 and 106 total cells). Chi-square analysis was used to compare groups. (D) Bar graph shows the percentage of mobile microglia for each mouse in each experimental group and condition. The number of mice per group is in parentheses. P values were derived form two-tailed Mann-Whitney tests. Data are mean ± SEM. (E) Linear regression analysis showing the relationship between absolute distance moved over a 24h period as a function of initial distance from the bleed in each experimental group. (F) Graphs show data for each cell and average absolute (top row) or net (bottom row) movement of mobile microglia in each experimental group under normal and injury conditions. Two way ANOVA did not reveal any main effects of IFNγ status or bleed. Data are mean ± SEM.

Given that sex strongly influenced microglia mobility in our previous injury experiments, we present data for male (**Fig. 3**) and female mice (**Fig. 4**), separately. In male mice without any injury, we did not find a single mobile microglia in mice with cre-recombinase dependent knockdown of *ifngr1* (9 mice, 122 cells imaged; **Fig. 3C,D**). The absence of mobility in male *ifngr1* KD mice contrasted with WT mice injected with IFNγ or vehicle/control protein (compare *ifngr1* KD with WT and WT + IFNγ “no bleed” data in **Fig. 3C,D**). Following the induction of a microbleed, *Ifngr1* KD mice had significantly fewer mobile microglia after injury (**Fig. 3B-D**), suggesting that their ability to migrate was impaired. By contrast, male mice injected with IFNγ protein exhibited significantly more mobile microglia than WT controls or *Ifngr1* KD mice (**Fig. 3B-D**). Plotting microglia movement as a function of distance from the CMB revealed that this relationship, which is of moderate strength and highly significant in WT control or WT mice injected with IFNγ protein, was lost in male *Ifngr1* KD mice (**Fig. 3E**). And finally, we plotted the absolute and net movement of all mobile microglia in each group (**Fig. 3F**). Our analysis did not indicate a significant main effect of Bleed, IFNγ status or interaction in any of the comparisons (all p values > 0.05). Thus, we conclude that in male mice, IFNγ signaling plays a role in mobilizing microglia under normal or injury conditions, but does not affect how far these mobile microglia move, or their directionality once activated.

**Figure 4.**
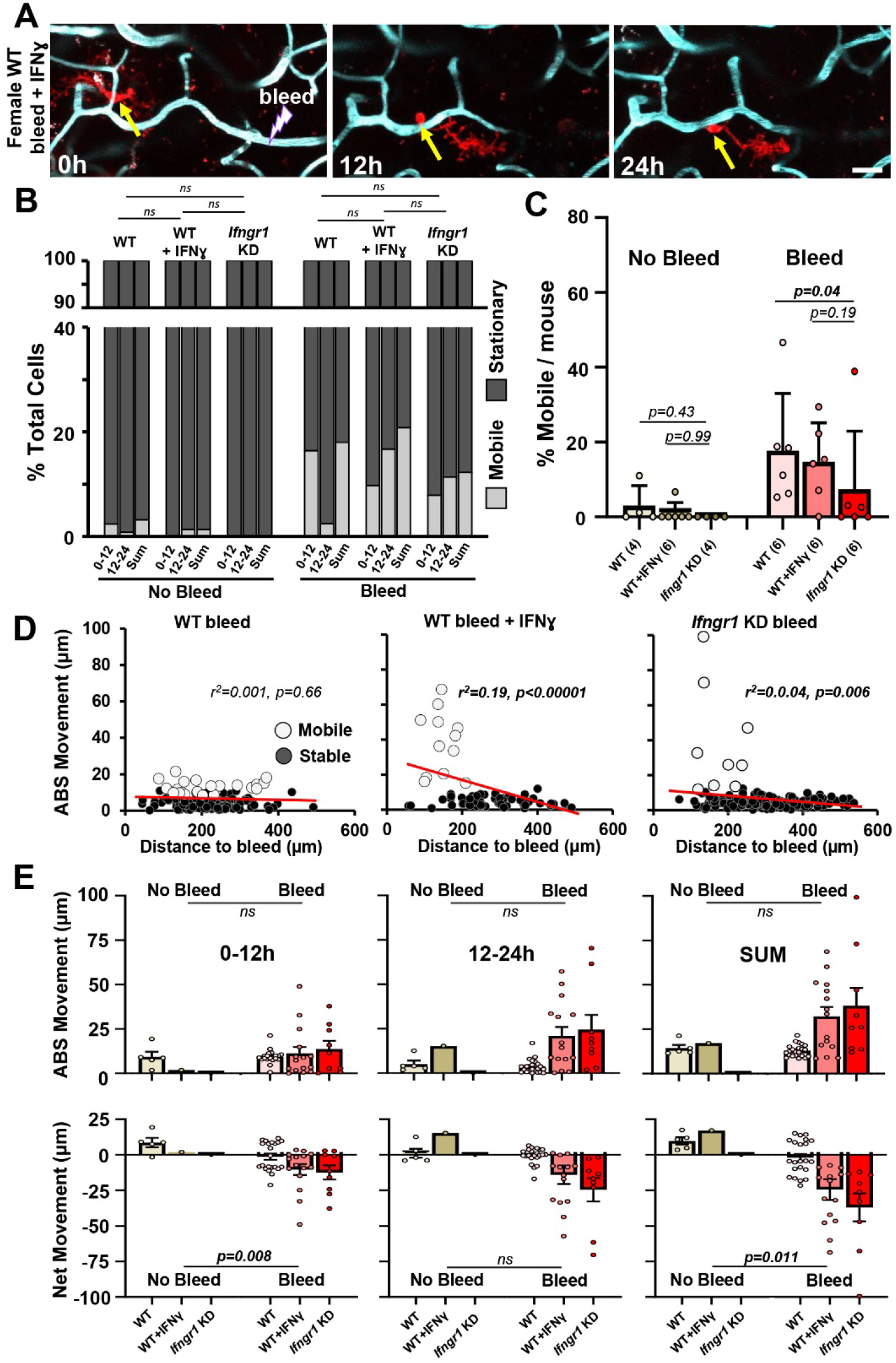
Minimal effect of IFNγ signaling on microglial mobility in female mice. (A) Two-photon images illustrate the migration of a microglia (yellow arrow) after injury in a female mouse stimulated with IFNγ. Scale bars, 20μm. (B) Fraction of mobile or stationary microglia in each experimental group or injury condition (WT no bleed and bleed: 153 and 122 total cells; WT+IFNγ no bleed and bleed: 75 and 72 total cells; IFNgR1 KD no bleed and bleed: 121 and 161 total cells). Chi-square analysis was used to compare experimental groups. (C) The percentage of mobile microglia per female mouse in each experimental group and injury condition. Number of mice per group are in parentheses. Between group comparisons were derived form two-tailed Mann-Whitney tests. Data are mean ± SEM (D) Linear regression illustrates the relationship between absolute movement over a 24h period and original distance from the injured vessel. (E) Graphs show data for each cell and the average absolute and net movement of mobile microglia in each experimental group and injury condition. Two way ANOVA followed by two-tailed unpaired t-tests were used to calculate p values. Data are mean ± SEM.

Next we examined whether IFNγ signaling regulates microglia mobility in female mice (**Fig. 4A**). In the absence of injury, we did not detect a single mobile microglia in female *Ifngr1* KD mice (n=4 mice, 121 cells; **Fig. 4B**). However, since female WT mice tended to have fewer mobile microglia in general, there were no significant group differences in the % mobile microglia as a fraction of total cells sampled or % cells per mouse (**Fig. 4B,C**). In contrast to our observations in male mice following injury, we did not find a significant increase in mobile microglia in female WT mice injected with IFNγ, nor did we find a significant decrease in the proportion of mobile microglia in *Ifngr1* KD mice (**Fig. 4B,C**). In contrast to our findings in males (**Fig. 3E**), the relationship between the initial distance from the bleed and absolute distance travelled after bleed, was absent or very weak in female WT and *Ifngr1* KD groups respectively (**Fig. 4D**; r^2^ values: 0.001-0.04), whereas a moderate relationship was evident in WT female mice stimulated with IFNγ (**Fig. 4D**; r^2^=0.19). Probing the absolute or net distance travelled by the small fraction of mobile microglia did not reveal any significant effects of IFNγ status or Bleed x IFNγ status interaction at any time point examined (all p values > 0.05, **Fig. 4E**). However, there was a significant main effect of bleed (vs No bleed) in net movement at 0-12h and the sum of movements over 24h (**Fig. 4E**). Thus, in contrast to our findings in male mice, IFNγ signaling appears to play a lesser role in regulating microglia mobility in female mice.

## Discussion

The extent to which microglia can mobilize in the mature mammalian cerebral cortex, has mostly been studied using histology or densely labelled transgenic mice. Furthermore, few studies have focused on the molecular mechanisms that regulate this behavior, at least *in vivo*. In the present study, we quantitatively assessed the mobility of individual microglia under normal and injury related conditions in mice with sparsely labelled microglia. Consistent with previous studies, our experiments show that only a small fraction of microglia are mobile under normal conditions. After injury, the fraction of mobile microglia increased significantly, with some cells moving an impressive 60-100μm in a 24h period, especially when situated close to the injury. However, the capacity for some microglia to mobilize was strongly influenced by sex. For example, microglia in male mice were more likely to mobilize after injury, moved significantly greater distances, and showed greater directionality in movement when compared to microglia in female mice. In order to further understand the factors that regulate mobility, we probed the role of the pro-inflammatory cytokine IFNγ, in both male and female mice. Our experiments revealed that IFNγ signaling plays an important role in promoting microglia mobility, but primarily in male mice. Collectively, these findings reveal the heterogenous capacity for migration in microglia and the powerful influence that sex and IFNγ signaling play in this unique cellular behavior.

In agreement with previous studies, only a small fraction of microglia are mobile under normal conditions^16–18^. However, the explanation for why so few mobilize remains speculative. Since microglia undergo cell death at low rates comparable to the percentage of cells that mobilize, it is conceivable that these cells move in order to re-populate territory vacated from dying or dead cells^16,17^. Dying cells would trigger the opening of NMDA receptors and release of chemoattractant signals like purines or lysophosphatidylcholine, which previous work has implicated in cell migration^16,54,55^. Purine release could also come from the stochastic opening of the blood brain barrier, such as microbleeds that do occur in healthy brains (insert reference). However, these rare events would seem unlikely to account for the ~5% of cells that move per day. Very recent work has shown that microglia are highly attracted to bouts of neural activity^56^. For example, increased neural activity can trigger NMDA receptor activity and purine release^57^, which stimulates microglia process envelopment and suppression of active neurons^58^. Whether similar mechanisms control whole body movement of microglia remains an open question for future studies.

Following the induction of a microbleed, we show increased mobilization of microglia that ultimately cluster around injury sites^14,59^. Of note, some cells could travel up to ~100μm within a 24h period, which is further than past studies, likely due to previous limitations noted with single cell tracking. This finding reinforces the idea that microglia retain a remarkable capacity for migration well into adulthood. Indeed, this has been noted in one previous study showing putative microglia can migrate away from an injury site (2-4 weeks after injury), on the order of millimeters^30^. Another finding was that the recruitment of microglia was strongly dependent on their initial distance from the site of injury, with the majority of mobile cells residing within 300μm^14^. This dependence on distance implies that a diffusible, injury related factor was contributing. Based on previous work examining microglial process chemotaxis, there are several plausible mechanisms. The release of purines from damaged cells next to the injury site^60^ or fibrinogen from blood plasma, are established signals for stimulating microglial process chemotaxis towards injury^26,29,60^. Microbleeds can also locally change neural activity, although usually this is associated with a suppression of activity^61^. Given that microglia are attracted to increased neural activity^56,58^, this would seem less likely to explain mobility towards the injury. Local changes in electrolytes or purines could also alter THIK-1 channel activity which regulates microglia transition states between surveillance and injury induced chemotaxis, as well as the release of pro-inflammatory cytokines^32^.

Gene expression studies have firmly established that microglia express IFNγ receptors^34^ and upregulate IFNγ response genes after injury or neurodegeneration^49^. Vascular insults such ischemia and hemorrhages rapidly mobilize IFNγ secreting leukocytes to sites of injury^14,62^. Pro-inflammatory cytokines like IFNγ can readily pass through the blood brain barrier^63^ and stimulate a reactive, amoeboid like microglial morphology, secretion of pro-inflammatory cytokines^36,37,64^ and migration, at least in cell culture^37^ and retina^33^. In the mature brain, our previous imaging study showed that IFNγ could affect microglia process chemotaxis towards vascular injury^50^. However, despite the likelihood that IFNγ signaling could regulate microglia migration, direct *in vivo* tests of this in the adult cortex were lacking. Here, we show that stimulating IFNγ signaling by doping the blood with recombinant IFNγ before microbleed, upregulates the fraction of mobile cells but does not significantly alter the absolute or net distance microglia travel. Conversely, genetic knockdown of *ifngr1* specifically within male microglia, blunted the fraction of microglia that could be mobilized, but not the overall distances they travel. These results suggest that microglia mobility is like a switch. Once they are stimulated to become mobile cells (by injury or IFNγ signaling), the distance with which they move is no longer tightly regulated by these stimulants. Even in the absence of injury, microglia in *ifngr1* knockdown were much less mobile, as we did not find a single mobile cell in 13 mice (9 male, 4 female). This finding suggests that IFNγ signaling may exert some tonic effect on mobility in the normal brain, perhaps though low level expression of IFNγ in resident immune cells, or conceivably constitutive IFNγ receptor activity. Consistent with this idea, the Kipnis group^51^ showed that IFNγ secreting cells in the meninges can regulate neural and social activity in healthy mice, presumably through diffusion of IFNγ in cerebrospinal fluid to neurons. In the context of the present work, immune cells that secrete IFNγ such as T cells and monocytes, which are normally present in the meninges or recruited to sites of injury^65^, could directly communicate with microglia to regulate mobility. Indeed, a fascinating study by^66^ showed that the recruitment of T cells after ischemic stroke, activated microglia in the peri-infarct region (i.e. de-ramified morphology) and induced gene expression associated with interferon signaling and chemotaxis. Whether IFNγ based mobilization of microglia is of help or hindrance to normal brain function or repair after injury, will require further investigation. Systemically stimulating IFNγ signaling after middle cerebral artery occlusion exacerbates ischemic damage^62^. Similarly, upregulation of IFNγ signaling in microglia after viral infection promoted synapse loss and cognitive deficits in mice^52^. While beyond the scope of the present study, future studies could manipulate IFNγ signaling to promote or limit microglial mobilization towards lesions associated with vascular injury or multiple sclerosis^67^, and determine if they hold any therapeutic benefit. This more subtle approach might be preferable to microglial depletion approaches which have the unintended consequence of facilitating the recruitment of potentially deleterious infiltrative macrophages^67–69^.

A common thread through most of our experimental results was the dependence on biological sex. Indeed, we found that microglia in male mice were significantly more likely to become mobile after injury than female mice, and these mobile cells travelled greater distances. Male microglia generally mobilized towards the microbleed whereas microglia in female mice showed less directionality, at least on average (Note that mean Net movement is close to 0 for females in Fig. 2F). Furthermore, stimulating or inhibiting IFNγ signaling had much greater effects on stimulating mobility in microglia from male mice than females. Together, these results suggest that microglia in male mice may be more primed to migrate in response to injury or inflammatory related signals than female mice. In recent years, there has been a greater appreciation for sex differences in microglia function and gene expression in the context of normal and disease related conditions^70–72^. For example, RNA sequencing^41^ showed that microglia from male mice were enriched with genes and proteins associated with inflammation and chemotaxis^38,41–43^. Microglia in adult male mice were also significantly larger and more branched, display greater ionic conductance at rest and stronger inward currents in response to ATP^41–43^. While sex differences in migratory abilities in the cortex have not been extensively studied *in vivo*^38^, *in vitro* data show that male microglia can migrate greater distances than female microglia, when stimulated with IFNγ. Similarly, male brains show significantly higher levels of IFNγ when stimulated with LPS than female brains^73^. These findings could explain in our study why microglia in male mice were much more affected by manipulations of IFNγ than female mice. At the very least it suggests that female microglia rely on additional factors to become mobile. Why these sex specific differences in response to microbleed or IFNγ exist, is unclear. For example, sex hormones can account for microglial differences in spinal hyper-excitability and pain sensitivity^71^, but less so for gene expression differences^41^. Sex chromosomes can also independently influence microglia function and gene/protein expression, and in fact strongly modulate sensitivity to stroke damage in aged mice^74^. Single cell analysis of the transcriptome in mobile cells in male and female mice would be particularly fascinating, albeit technically challenging since only a subset of microglia show mobility, and these would need to be first identified *in vivo*. Nonetheless, our results should stimulate future studies examining the influence of sex hormones, chromosomes and differential gene expression on microglia mobility.

## Methods

### Animals

Adult male and female mice between 2 and 6 months of age were used in this study. For labelling of microglia and monocyte derived macrophages, experiments involved heterozygous CX3CR1^+/GFP^ mice (JAX# 005582) on a C57BL/6J background (JAX# 000664). For cre-dependent inducible expression of the tdTomato reporter and/or knockdown of *Ifngr1*, we utilized Ai9 reporter mice (B6.Cg-*Gt(ROSA)26Sor^tm9(CAG-tdTomato)Hze^*/J, JAX# 007909) crossed with an inducible microglia specific cre driver line (*Tmem119^em1(cre/ERT2)Gfng^*/J, JAX# 031820). These Ai9:Tmem119^CreERT2^ mice were then crossed with floxed *Ifngr1* mice (C57BL/6N-*Ifngr1^tm1.1Rds^*/J, JAX# 025394) and bred to homozygosity. All mice were housed in groups on a 12h light/dark cycle in ventilated racks in a humidity (RH 40-55%) and temperature controlled room (21-23°C). Mice were provided food and water *ad libitum*. All experiments comply with the guidelines set by the Canadian Council on Animal Care and approved by the local university Animal Care Committee. Reporting of this work complies with ARRIVE guidelines.

### Surgical Procedures

Mice of at least 2 months of age, were anesthetized using isoflurane (2% for induction and 1.3% for maintenance) in medical air (80% N2, 20% O2) at a flow rate of 0.7L/min. A temperature feedback regulator and rectal probe thermometer maintained animals’ body temperature at 37°C throughout the procedure. After subcutaneous injection of lidocaine under the scalp, an incision was made along the midline. A custom metal ring (~1g in weight, outer diameter 11.3mm, inner diameter 7.0mm, height 1.5mm) was positioned over the right somatosensory cortex and secured to the skull with metabond adhesive. A circular area (diameter of ~4-5mm) of skull within the metal ring was thinned using a high-speed dental drill, and ice-cold HEPES buffered artificial cerebrospinal fluid (ACSF) was periodically applied to the skull for cooling purposes. Fine forceps were used to remove the thinned piece of skull and the exposed brain was kept moist with gel foam soaked in cold ACSF. A 5 or 6mm coverslip (no. 1 thickness) was placed over the exposed brain and secured to the skull using cyanoacrylate adhesive. After the procedure, mice were injected with 0.03mL of 2% dexamethasone (i.p.) to reduce acute inflammation resulting from the procedure. Mice were monitored while they recovered under a heat lamp and then were returned to their home cage.

### Experimental treatments and *in vivo* two-photon imaging

Two to four weeks after cranial window implantation, cre-recombinase dependent expression of tdTomato reporter and/or knockdown of *Ifngr1* was induced with an intraperitoneal injection of 0.5mg of Tamoxifen (Sigma #T5648) dissolved in corn oil (Sigma #T5648). If mice failed to show sufficient reporter expression two weeks later, they were reinjected with the same dose of tamoxifen.

For two-photon imaging, mice were lightly anesthetized with isoflurane (2% for induction and 1% for maintenance in medical air), and then received 0.1mL of either 1.5-3% Fluorescein isothiocyanate–dextran (FITC) or Texas Red dextran (70kDa, Sigma-Aldrich #46945 and Thermofisher D1830, respectively), to permit visualization of the cerebral vasculature. The metal ring on the mouse’s head was fixed into a custom imaging stage. High-resolution two-photon image stacks of the vasculature and GFP or tdTomato expressing microglia/macrophages were acquired *in vivo* using an Olympus FV1000MPE laser scanning microscope fed by a mode-locked Ti: Sapphire laser source (Mai Tai XF DeepSee, Spectra-Physics) and equipped with a water-dipping 20× objective lens [Olympus; numerical aperture (NA) = 0.95]. The laser was tuned to 900nm for imaging Texas red dextran and eGFP, while 945nm was used for imaging FITC and tdTomato labelled microglia. Emitted light was split by a dichroic filter (552 nm) before it was directed through band-pass filters (495 to 540nm and 558 to 630nm). Images were collected at 1.75-μm z-step intervals from the cortical surface to a depth of 75-200μm, with each image covering an area of 635.3 x 635.3μm (1024 x 1024 pixel sampling). In order to maximize sampling of sparsely labelled microglia, we typically imaged 2-5 different regions within each mouse, spaced at least 500μm apart from each other. Capillaries that were 3-6μm in diameter were targeted for laser induced rupture. As previously described, we raster scanned a 4×4μm region of interest centered on the capillary for 3-8s using the femtosecond laser (805nm, ~390mW at back aperture, 10μs pixel dwell time). The rupture of a cortical capillary was confirmed in real-time by the extravasation of fluorescently labelled blood plasma. Brightfield images of the brain’s surface and vascular landmarks were used to re-locate imaging areas over time.

For modulating IFNγ signaling *in vivo*, we used two approaches. First for stimulating IFNγ signaling^62^, we randomly assigned Ai9:Tmem119cre mice to receive an intravenous injection of control solution or 0.05 mg/kg of recombinant IFNγ from mouse (Sigma #I4777), minutes before initiating the rupture of cortical capillaries and then 12h later. Control solution injections consisted of injecting vehicle with or protein control (albumin or isotype control antibody, clone 2A3, Bio X Cell) or albumin. Our analysis indicated that there were no significant differences in the fraction of mobile microglia or ABS migration distance between vehicle and control protein injected mice (% mobile cells: 2 tailed t-test, t_(5)_=0.75, p=0.48; ABS movement Sum: 2 tailed t-test, t_(30)_=0.89, p=0.38), therefore data from these groups were pooled together under “WT bleed” group.

### Data analysis

In order to estimate error associated with 3-dimensional measurements *in vivo*, we repeatedly imaged GFP labelled VIP neurons in mice (*Vip^tm1(cre)Zjh^*/J, JAX# 010908) at 12h intervals (see Supp. Figure 1). Using Neurolucida software, we quantified the Euclidean distance from the center of each cell body to a fiducial vascular landmark in the center of the image. Performing this analysis in 3 mice with 206 neurons, yielded a mean error of 1.84±1.45μm and 1.61±1.53μm for 0-12 and 12-24h measurements. To minimize any false positive errors, we set the minimum migration cut-off at 4 standard deviations above the mean error values to yield a minimum cut-off migration distance of 7.46μm in a 12h period.

Having established our criteria for detecting true migration, we then classified microglia as “mobile” or “stationary” based on whether their 3D position from a fiducial vascular landmark (typically a branch point) changed by more than 7.46μm at 0-12 or 12-24h. The sum of mobile cells represented all those that exceeded this criteria at either or both time intervals. Since cells could move towards or away from a vascular landmark or bleed site, movement was calculated as either the absolute value (thus indifferent to direction) or the net value relative to vascular landmark or bleed location. For the analysis of movement in capillary associated microglia (CAM), we first determined whether the closest edge of a microglia soma was within 1.86μm (3 pixels) from the lumen of a flowing a capillary or not. Based on this dichotomy, we than analyzed the absolute and net movement of each microglia across and within sexes. In order to estimate the extent of plasma extravasation after microbleed, we calculated extravasation area from single images collected immediately after the rupture that were then mean filtered (radius=2 pixels) and thresholded using Huang’s method in Image J.

### Statistics

Absolute and net distances travelled by mobile microglia were analyzed with two-way ANOVA using Time, Sex, injury (bleed or no bleed) and IFNγ status (WT control vs WT+ IFNγ vs *Ifngr1* KD) as factors. Post hoc tests were adjusted using Sidak’s multiple comparisons tests. Chi-squared analysis was used to determine if the fraction of mobile cells differed between experimental groups. The “observed” values for each experimental group were compared to the “expected” values, which were derived from No bleed or Male groups in Figure 2, or WT group in Figures 3 and 4. Linear regression analyses were used to determine the relationship between a cell’s initial distance from a microbleed and the absolute distance it travelled over 24h after bleed. Non-parametric tests (2-tailed Mann-Whitney test) were used to compare groups when examining the % mobile cell per mouse. Unless otherwise stated, data are presented as mean ± SEM. Cut-offs for significant differences were as follows: *p<0.05, **p<0.01, ***p<0.001.

## Acknowledgements

We are grateful to Manu Rangachari and Patrick Reeson for their advice and assistance with the project, as well as Angie Hentze and Taimei Yang for managing the mouse colony. Work was supported by operating, salary and equipment grants to C.E.B. from the Canadian Institutes of Health Research (CIHR), Heart and Stroke Foundation (HSF), Natural Sciences and Engineering Research Council (NSERC).

## Author Contributions

R.B performed the experiments, analyzed the data and wrote portions of the manuscript. S.S. and M.C. contributed to data collection and analysis. C.E.B conceived the study, contributed to data collection, analysis and wrote the manuscript.

## Data Availability

Data generated for this study are available from the corresponding author reasonable request.

