## Supplementary data for "Sex and interferon gamma signaling regulate microglia migration in adult mouse cortex *in vivo*"

### Supplementary Figure 1

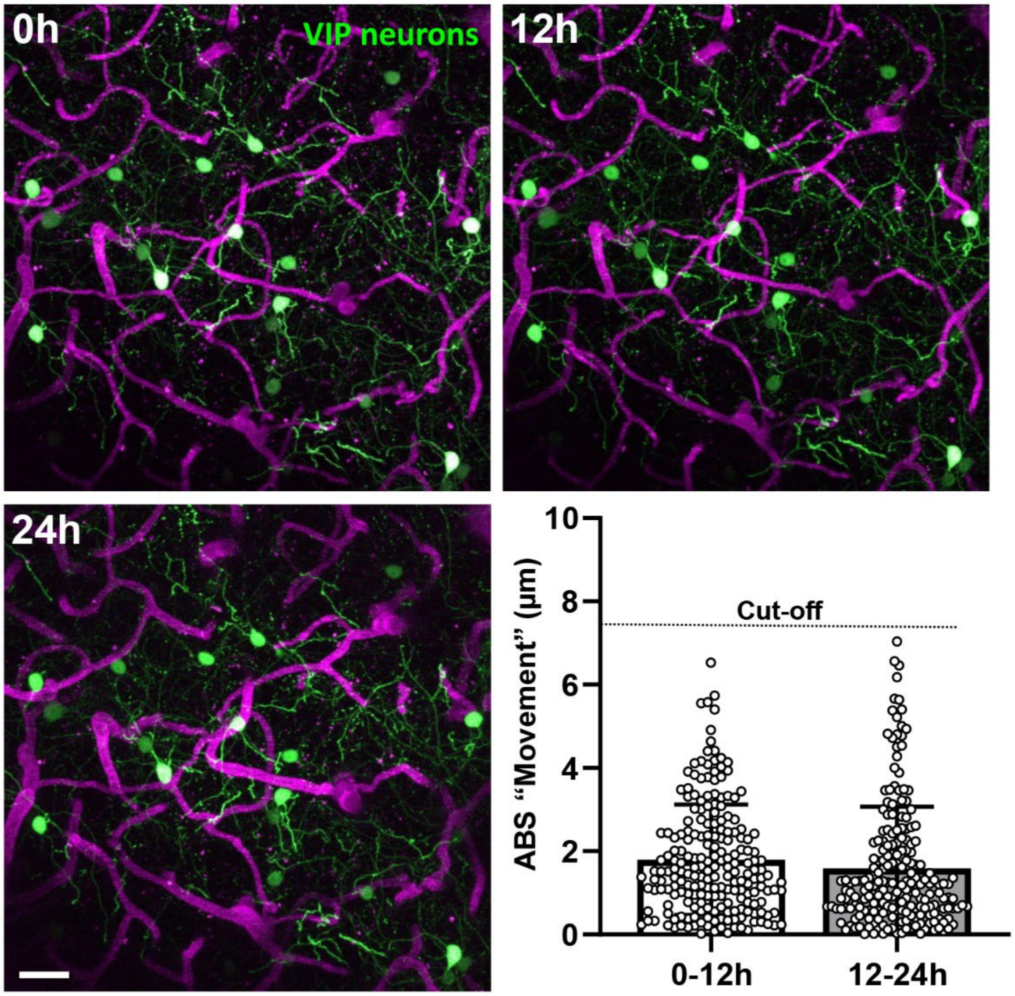


**Supplementary Figure 1. Approach for defining "true" cell movement *in vivo.*** Two-photon *in vivo* images show GFP labelled VIP neurons and cortical vasculature imaged at 12h internals over 24h. The absolute distance of each neural cell body to an arbitrary landmark (eg. Vessel branch point) was measured to calculate a mean error value. The cut-off point for defining a cell as "mobile" was 4SD above the mean. Data are mean ± SEM. Scale bar, 20μm.

### Supplementary Figure 2

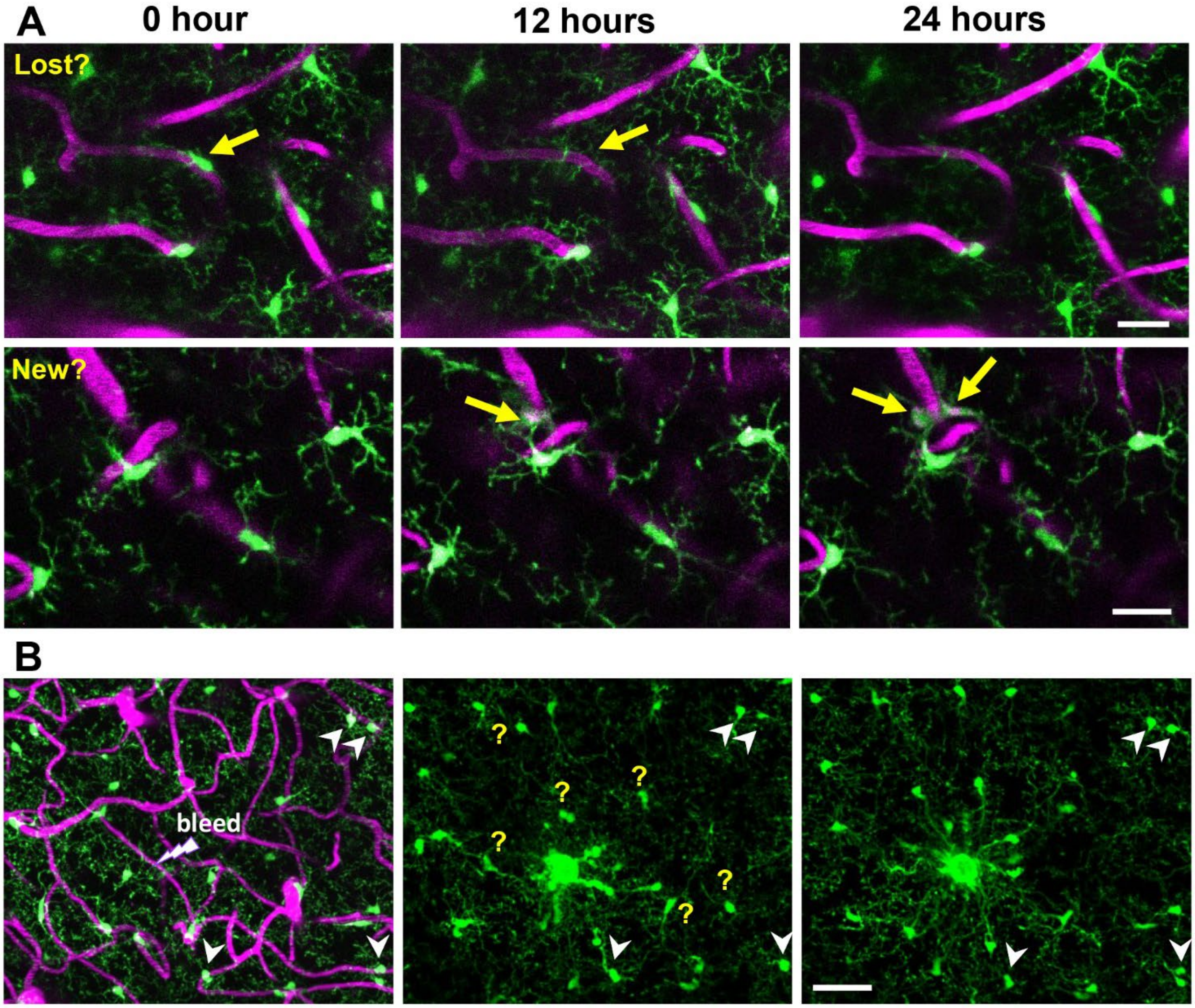


**Supplementary Figure 2. Time lapse imaging of microglia movements under normal conditions in a Cx3cr1-GFP mouse.** (**A**) Two-photon z-projection images show Cx3cr1-eGFP labelled microglia and cortical vasculature imaged at 12h internals over 24h. We noted the occasional instance of microglia loss (top row) or appearance of a new cell (bottom row). Scale bar, 20µm. (**B**): Two-photon image projections showing re-arrangement of Cx3cr1-eGFP microglia/macrophages over 12h intervals after the induction of a microbleed. Note that some cells distant to the bleed appear stable (white arrowhead). However, many cells appear in new positions or disappear (perhaps into the center of the injury site), making single cell tracking untenable. Scale bar, 50µm. Data are mean ± SEM.

### Supplementary Figure 3

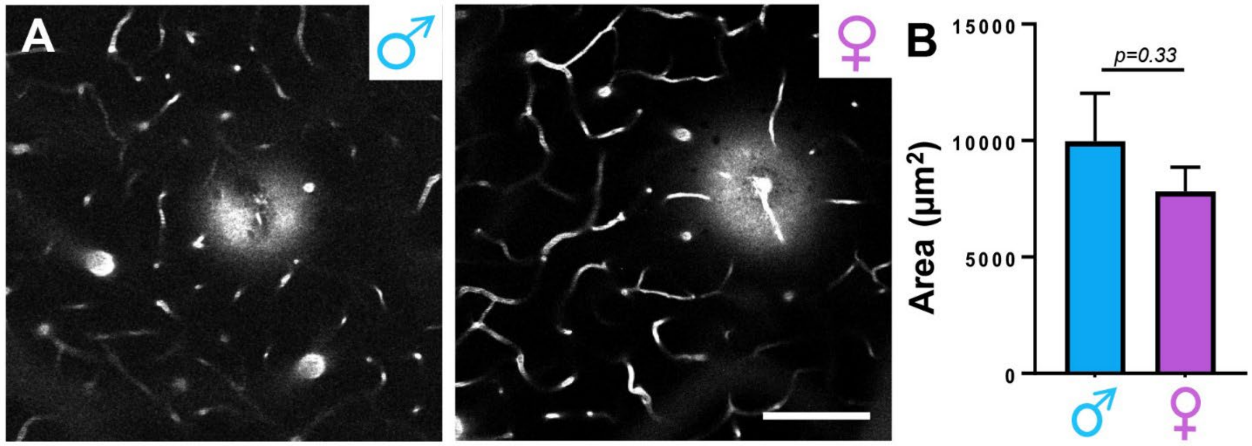


**Supplementary Figure 3. Estimation of bleed size based on extravasation of fluorescent plasma.** (**A**): Single plane *in vivo* images show extravasation of FITC labelled blood plasma immediately after rupture of a single capillary in a male and female mouse. Scale bar, 100µm. (**B**): There was no sex difference in the area of dye extravasation. Two-tailed unpaired Welch's t-tests were used to calculate p values. Data are mean ± SEM.

### Supplementary Figure 4

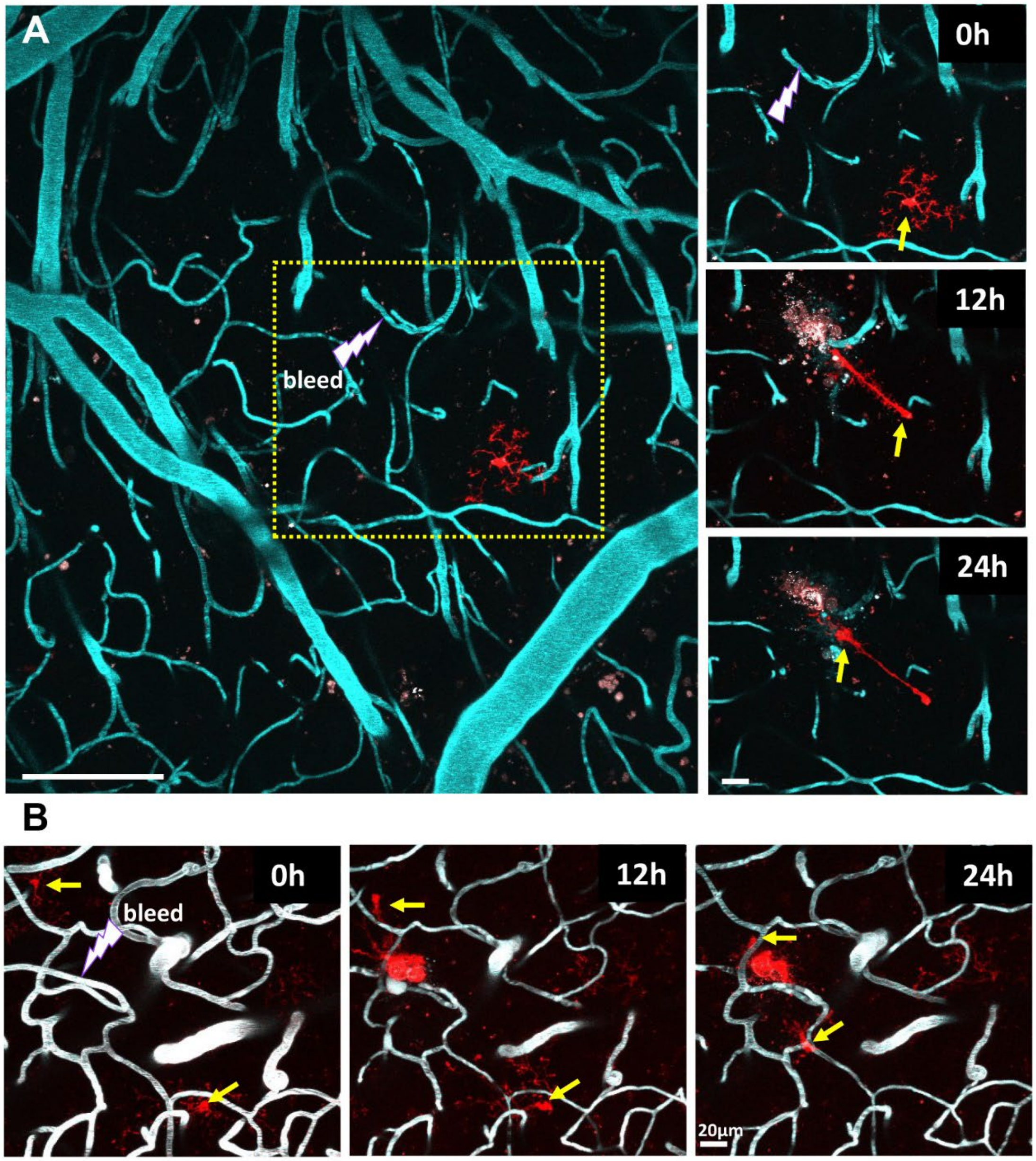


**Supplementary Figure 4. Examples of microglia migration in male mice after vessel injury.** (**A**): *In vivo* images show tdTomato labelled microglia (red) and cortical vasculature in a male mouse before the rupture of a capillary and then 12 and 24h later. Scale bar for left panel, 100µm; right panel, 20µm. (**B**): Example of two microglia migrating towards the injury site in a male mouse treated with IFNg. Yellow arrows denote migrating cells. Scale bar, 20µm.

### Supplementary Figure 5

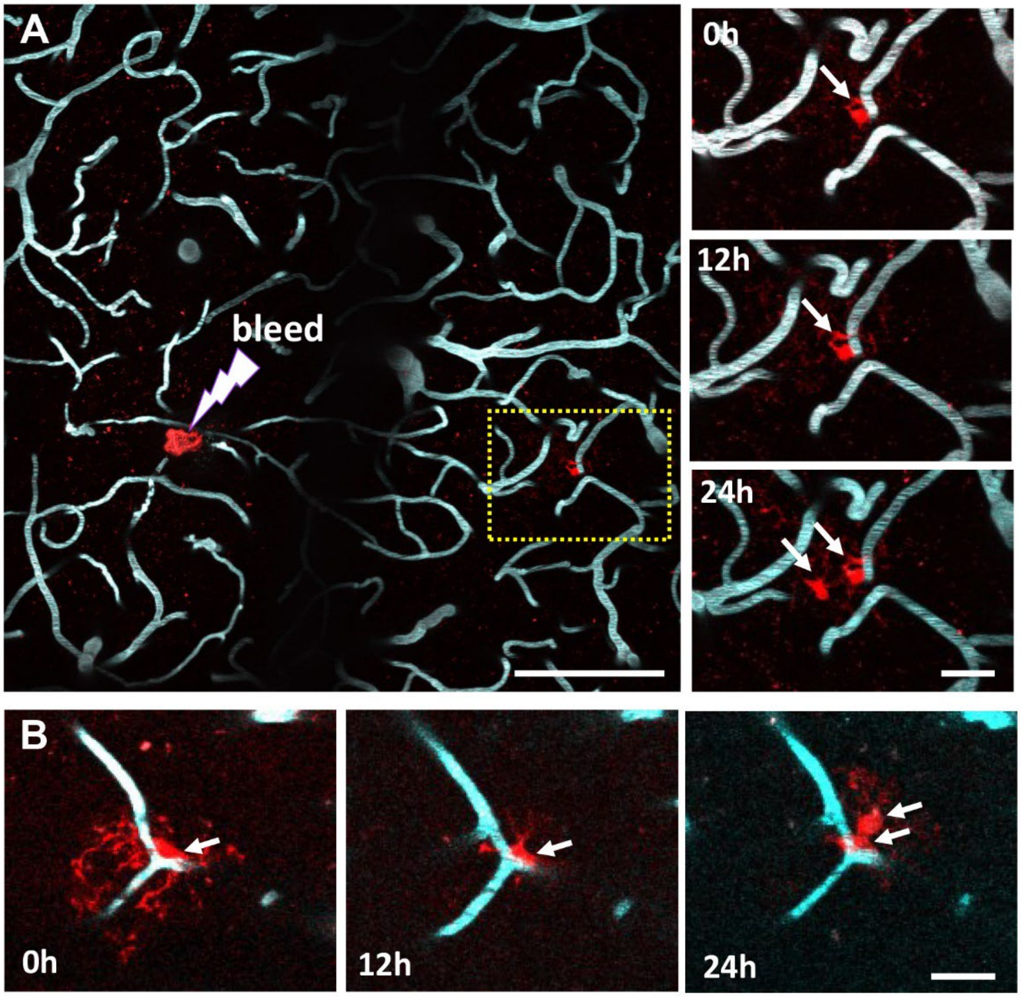


**Supplementary Figure 5. Microglia proliferation after injury.** (**A**): *In vivo* images show tdTomato labelled microglia (red) and cortical vasculature in a male mouse before and after vessel injury. The new cell appears at the 24h time point. Scale bar for left panel, 100µm; right panel, 20µm. (**B**): Images show microglia proliferation in a male *Ifngr1* KD mouse. Scale bar, 20µm.

### Supplementary Figure 6

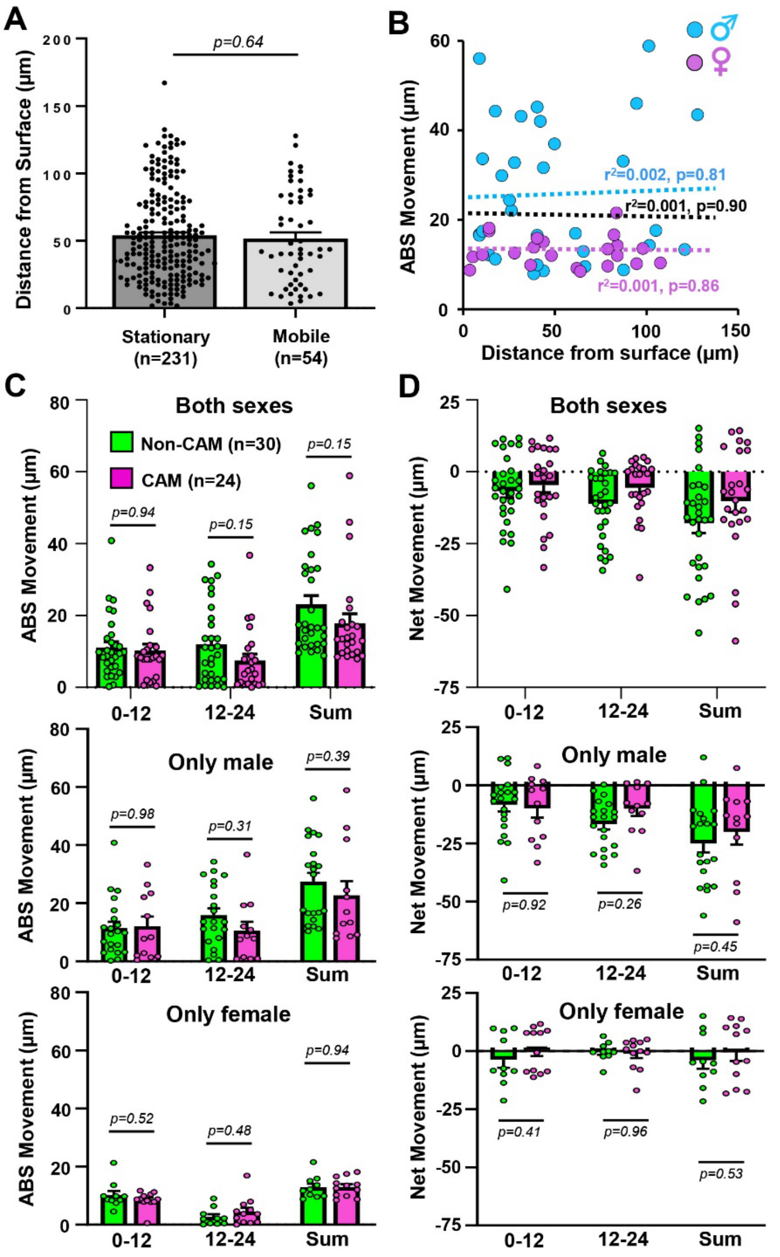


**Supplementary Figure 6. Microglia movement after injury is not predicted by its depth from the cortical surface or whether it was attached to a capillary.** (**A**): Bar graph shows the depth from the cortical surface for microglia defined as stationary or mobile. Two-tailed unpaired t-test was used to calculate p values. (**B**): Scatterplot of mobile microglia shows there is no relationship between the absolute distance travelled over 24h and how far below the cortical surface each cell resided, regardless of sex. Statistical values were derived from linear regression analysis. (**C**): Graphs indicate the absolute distance travelled over 0-12h, 12-24h or the sum of both time periods in mobile microglia associated with a capillary (“CAM”) or those that were not (“Non-CAM”). (**D**) Graphs reveal the net distance travelled by mobile CAM and non-CAM. Data in C and D are shown with both sexes pooled (top row), or just male (top row) or female (bottom row) mice. Two-tailed unpaired t-tests were used to calculate p values. Data are mean ± SEM.
